# Multi-virulence and phenotypic spread of *Campylobacter jejuni* carried by chicken meat in Brazil

**DOI:** 10.1101/2021.12.14.472542

**Authors:** Phelipe Augusto Borba Martins Peres, Roberta Torres de Melo, Paulo Marcel Armendaris, Fabiano Barreto, Tiago Follmann Perin, Ana Laura Grazziotin, Guilherme Paz Monteiro, Eliane Pereira Mendonça, Eduarda Cristina Alves Lourenzatto, Arthur Slompo Muniz Bicalho, Marcelo de Vito Filho, Ana Beatriz Garcez Buiatte, Daise Aparecida Rossi

## Abstract

*Campylobacter jejuni* is the most incriminated pathogen in bacterial gastroenteritis, and therefore, characteristics of its epidemiology must be continuously investigated to support possible mitigating measures. This is particularly important when evaluating representative strains of the world’s leading chicken meat exporter, Brazil. We evaluated a panel of 14 virulence genes in 359 strains of *C. jejuni* isolated from chilled broiler carcasses of Brazil. The genes were classified into five virulence categories (B: biofilm/motility; SS: secretion/cytotoxicity system; CI: invasion/colonization; GB: Guillain-Barré and AE: adaptation to stress). The percentage of strains with stress adaptation genes (86.07%) indicates the potential to adapt to unfavorable environmental conditions and *hcp* gene in 97.77%, indicates the ability to cause serious infections in humans. Genes related to GBS in 77.44% of strains are an additional concern, which must be monitored. The gene panel showed the presence of 124 virulence profiles. Individual analyzes by carcass, slaughter establishment, and municipalities where they were located showed high I.Var., of 0.82, 0.87 and 0.78, respectively. Georeferencing indicated state A as a hotspot for virulent strains. Higher levels of isolation and multi-virulence were identified in the summer, which in Brazil is hot and humid. Proteomics was able to discriminate the strains, but due to the high heterogeneity between them, it did not allow to explain their dissemination. Together, our results showed that the studied strains are a potential danger to public health and that there is an urgent need for their surveillance and the adoption of control measures, especially in state A.

**AUTHOR SUMMARY:** *Campylobacter jejuni* is a bacterium considered one of the main causes of foodborne illnesses and the consumption of undercooked chicken meat is one of the main sources of human infection. In Brazil, epidemiological studies of this pathogen are still scarce, when compared to countries with structured surveillance, as well as, its analysis is not required by public health agencies in any group of foods intended for human consumption. Here we investigate the epidemiology of C. jejuni strains isolated from chilled chicken carcasses in Brazil, determining virulent and multivirulent strains, by the origin of the sample and its phenotypic patterns. The strains showed a high potential for adaptation to the environment, being classified as virulent and multivirulent, with a seasonal pattern in the hottest and humid periods of the year. In state A, the strains with the highest evolutionary level were isolated, when compared to the other states in the region. We hope that this study will help to better understand the potential risks that C. jejuni poses to the population and support surveillance agencies in tracking and adopting measures to minimize the dangers that this pathogen poses to public health.

## INTRODUCTION

*Campylobacter jejuni* is one of the most frequent causes of bacterial gastroenteritis worldwide, being the most prevalent agent in countries with structured surveillance systems [1, 2]. In underdeveloped or emerging countries, however, there are problems in underreporting human infections and identifying positive foods. These facts, added to the multiplicity of reservoirs and transmission routes, geographic and seasonal distribution, hinder the complete understanding of its epidemiology [3].

This microorganism is considered fastidious, with a high potential to form biofilms, easily acquires the viable non-cultivable form under conditions of extreme stress, and has a variety of virulence factors. These genetic determinants guarantee not only the efficiency in the infectious processes in the human host but also the survival in hostile environments and are fundamental in the characterization of the different strains [4].

Some genes have been recognized for their importance in the genus *Campylobacter*, including *flaA* and *luxS* involved in the formation of biofilms, *cdtABC* and *hcp* related to secretion systems, *cadF, ciaB* and *pldA* related to invasion and colonization, *dnaJ, htrA*, and *cbrA* involved in adaptation to environmental stress and *neuA* and *cstII* related to the induction of Guillain-Barré Syndrome [5, 6, 7, 8].

Given this, conducting epidemiological analyzes becomes a difficult and at the same time fundamental task in adopting intervention measures for its control and prevention [9]. These analyzes are particularly relevant in the poultry industry since the consumption of chicken meat contaminated with *Campylobacter* is one of the main sources of infection for humans and Brazil is the world’s largest exporter of this food [10]. The aim of this study was to determine the virulent genetic potential and phenotypic spread of *C. jejuni* isolated from broiler carcasses destined for internal and external trade. In this study it was possible to develop an index of classification of virulent and multivirulent strains, aiming to assist further epidemiological studies of the species. One state of Brazil deserves attention for presenting characteristics of a hotspot where strains with the highest converging virulence and multivirulence rates are observed in the territory, demonstrating risk to the population.

## RESULTS

### Genes, virulence profiles, isolate analysis by sample

Fourteen genes associated with virulence were researched in all 359 strains of *C. jejuni* isolated from chicken carcasses. The evaluation by virulence categories found that the concomitant frequency of genes linked to adaptation to stress was significantly higher (86.07% - 309/359) compared to the remaining categories, and those related to the secretion were the least frequent (131/359 - 36.49%) (Table 1). The significant majority of the isolates (292/359 - 81.34%) had at least eight of the 14 studied genes (p <0.05 - Fischer test), which portrayed the high percentage of virulent strains 72.14% (229/359), of which 47.60% (109/229) were classified as multi-virulent. We found a total of 124 distinct virulence profiles, of which five comprised 40.1% (144/359) of the strains, with 76 strains belonging to a single multi-virulent profile. This fact determined the low variability index (I.Var. = 0.35) considering the 359 strains.

**Table 1.**
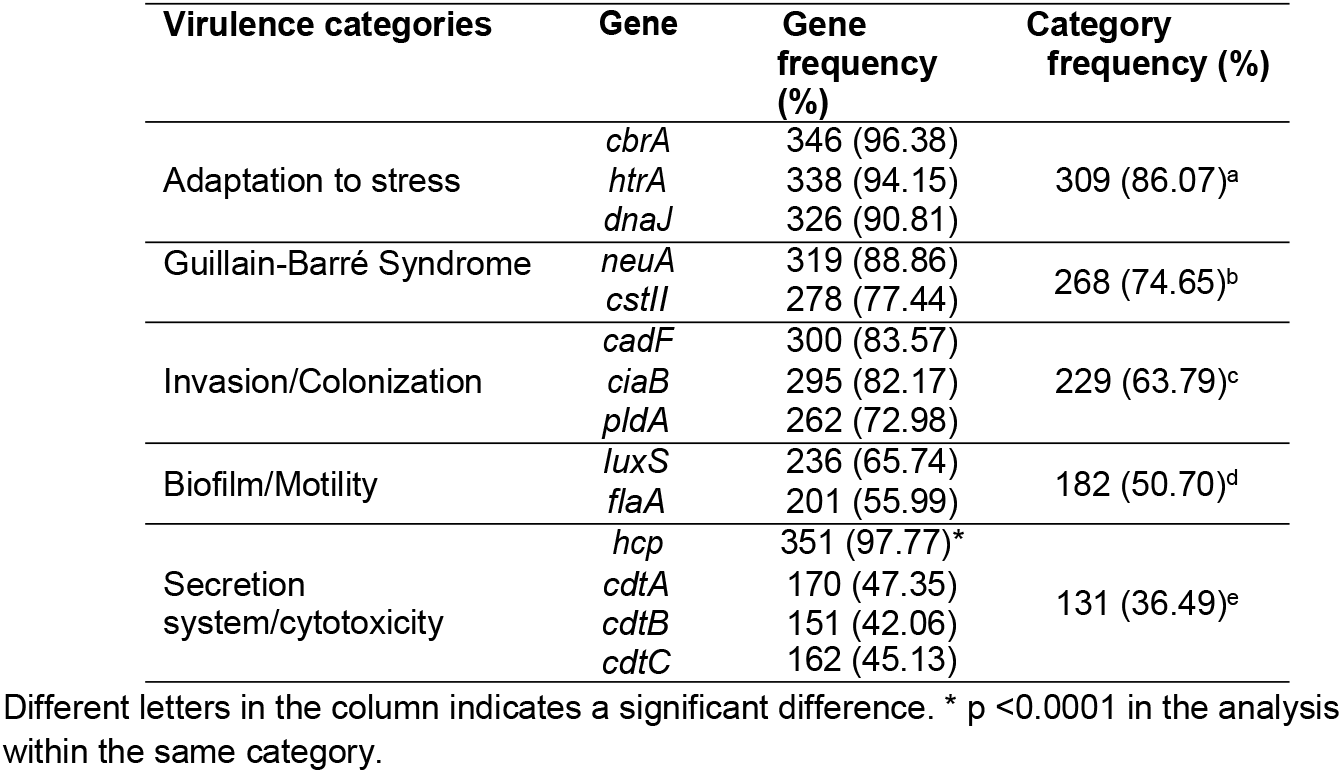
Frequency of virulence genes in *C. jejuni* isolated from chicken carcasses in Brazil between October 2017 and July 2018.

Profiles including genes linked to the biofilm formation/motility category were the least identified (60.5% - 75/124) and the ones associated with secretion systems (95.2% - 118/124) and adaptation to stress (96.0% - 119/124) were the most prevalent. The classification of the profiles according to the virulence categories allowed the identification of 16 clusters (A1 to A16), with A16 identified in 57 profiles (46.0%), which include virulent strains, 14 of which are profiles of multi-virulent strains (Table 2).

**Table 2.** Virulence profiles classified according to the categories of studied genes in *C. jejuni* isolated from chicken carcasses in Brazil between October 2017 and July 2018.

| Category | Number of profiles |
| --- | --- |
| Biofilm (B) | 75<sup>c</sup> (60.5 %) |
| Secretion system (SS) | 118<sup>b</sup> (95.2 %) |
| Invasion/Colonization (IC) | 106<sup>a</sup> (85.5 %) |
| Guillain-Barré Syndrome (GB) | 102<sup>a</sup> (82.3%) |
| Adaption to stress (AE) | 119<sup>b</sup> (96.0%) |
| Grouping | Number of profiles |
| A1. SS | 2 (1.61%) |
| A2. GB | 1 (0.80%) |
| A3. B, AE | 1 (0.80%) |
| A4. GB, AE | 1 (0.80%) |
| A5. IC, AE | 1 (0.80%) |
| A6. SS, AE | 3 (2.41%) |
| A7. B, GB, AE | 1 (0.80%) |
| A8. B, SS, IC | 1 (0.80%) |
| A9. B, SS, GB | 1 (0.80%) |
| A10. B, IC, AE | 1 (0.80%) |
| A11. SS, IC, AE | 2 (1.61%) |
| A12. SS, GB, AE | 6 (4.83%) |
| A13. B, SS, GB, AE | 2 (1.61%) |
| A14. B, SS, IC, AE | 11 (8.87%) |
| A15. SS, IC, GB, AE | 33 (26.61%) |
| **A16. B, SS, IC, GB, AE (Multi-virulent)** | **57\* (46.0%)** |
| **Total** | **124 (100%)** |

A: grouping. Different letters in the column indicates a significant difference. \*significant difference with the others. $p < 0.05$ . GB = Guillain-Barré Syndrome, AE = Adaptation to stress, CI = Cell invasion/colonization, SS = Secretion system/cytotoxicity, B = Biofilm/motility

The 359 strains used in our study came from the analysis of 114 carcasses, from which one to ten distinct colonies of *C. jejuni* were isolated per sample (Md = 3). Samples in which we obtained isolation from three or more colonies (59/114 - 51.75%) were grouped according to the number of isolates to determine the variability index (I.Var.), whose value identified did not differ between groups (p = 0.4841). The elevated I.Var. average found (0.82) justified the individual analysis of each strain throughout the study. The I.Var. identified within each group differed significantly (p <0.05), and for all samples, this value was greater than 0.75 (Figure 1).

**Figure 1.**
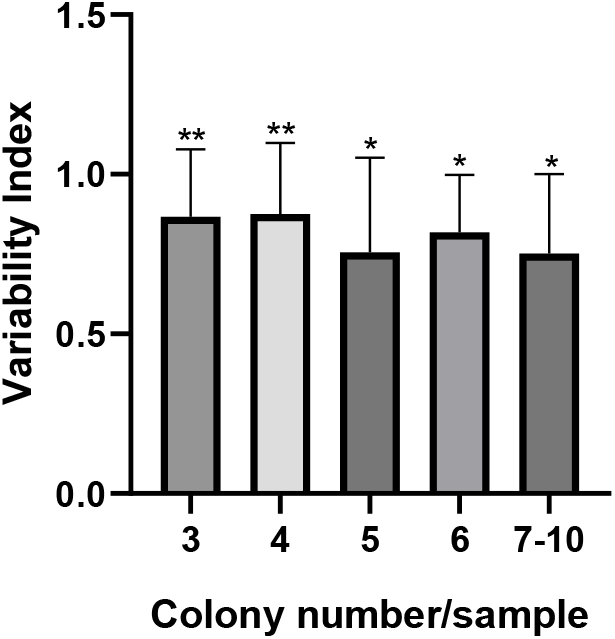
Variability index of *C. jejuni* per sample according to the number of isolated colonies. The error bar indicates the standard deviation, * p <0.05; ** p <0.01 by the Kruskall-Wallis test.

Our study included the analysis of 46 slaughterhouses under federal supervision in 43 municipalities. We obtained a variation of one to 22 isolates in each location, and those establishments that presented three or more strains (36/46 –78.3%) were used to determine the general and individual I.Var. scores. In the interpretation of virulence and multi-virulence indices, all establishments were considered.

The average I.Var. scores of all establishments was 0.87, with a significant majority of them (28/36 - 77.78%) showing values above 0.7 (Table 3).

**Table 3.** Variability index of virulence profiles of *C. jejuni* isolated from chicken carcasses by slaughterhouse.

| I.Var. of virulence profiles | Classification of I.V. | Establishments n=36 | Total |
| --- | --- | --- | --- |
| 1 | High | 10 (27.78%)<sup>ab</sup> | 28 (77.78%)<sup>a</sup> |
| 0.99-0.7 | High | 18 (50.00%)<sup>a</sup> |  |
| 0.69-0.5 | Medium | 7 (19.44%)<sup>bc</sup> | 8 (22.22%)<sup>b</sup> |
| <0.5 | Low | 1 (2.78%)<sup>c</sup> |  |

Different letters in the same column indicates a significant difference by Fisher's exact test ( $p < 0.05$ ).

The distribution of establishments according to the classification of virulence and multi-virulence is described in Table 4. In general and according to our expectations, the number of virulent strains per establishment and between establishments was significantly higher than that of multi-virulent strains (Student’s t-test - p <0.001). The significant majority of establishments (21/46 - 45.7%) had 70 to 100% of the strains classified as virulent, and two of them had the same percentage range of multi-virulent strains, both located in state A. Percentages less than 40% of virulent and multi-virulent strains were identified in 11/46 (23.9%) and 33/46 (71.8%) of the establishments, respectively.

**Table 4.**
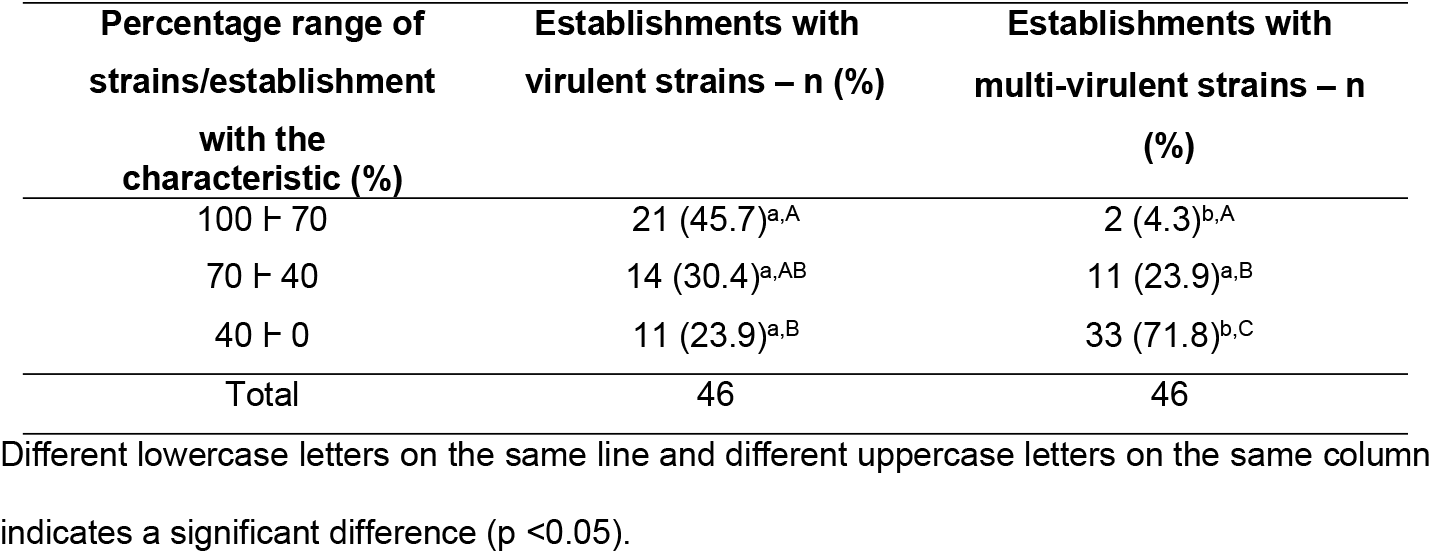
Frequency distribution of virulent and multi-virulent strains by establishments.

### Analysis of isolates by state, municipality and sazonality

The characteristics of *C. jejuni* were analyzed according to their states of origin: A, B and C. State A had the lowest number of isolates (81/359 - 22.6%) and profiles (42/124 - 33.9 %) compared to the other two states (p <0.05), whose numbers were similar between them (Table 5).

**Table 5.**
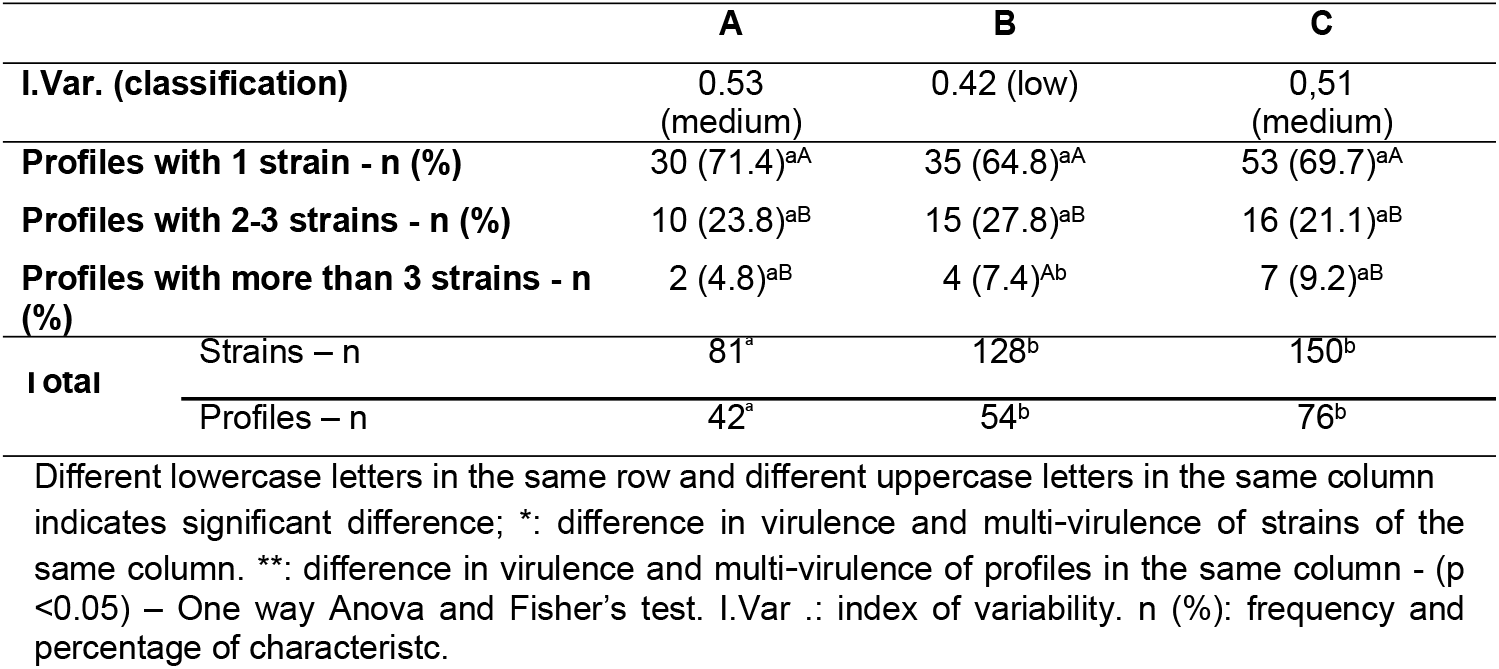
Variability indices, strain frequencies and virulence profiles identified in *C. jejuni* isolated from chicken carcasses in Brazil.

|  | A | B | C |
| --- | --- | --- | --- |
| I.Var. (classification) | 0.53 (medium) | 0.42 (low) | 0.51 (medium) |
| Profiles with 1 strain - n (%) | 30 (71.4)<sup>aA</sup> | 35 (64.8)<sup>aA</sup> | 53 (69.7)<sup>aA</sup> |
| Profiles with 2-3 strains - n (%) | 10 (23.8)<sup>aB</sup> | 15 (27.8)<sup>aB</sup> | 16 (21.1)<sup>aB</sup> |
| Profiles with more than 3 strains - n (%) | 2 (4.8)<sup>aB</sup> | 4 (7.4)<sup>Ab</sup> | 7 (9.2)<sup>aB</sup> |
| Total |  |  |  |
| Strains – n | 81<sup>a</sup> | 128<sup>b</sup> | 150<sup>b</sup> |
| Profiles – n | 42<sup>a</sup> | 54<sup>b</sup> | 76<sup>b</sup> |

Different lowercase letters in the same row and different uppercase letters in the same column indicates significant difference; \*: difference in virulence and multi-virulence of strains of the same column. \*\*: difference in virulence and multi-virulence of profiles in the same column - ( $p < 0.05$ ) – One way Anova and Fisher's test. I.Var.: index of variability. n (%): frequency and percentage of characteristic.

Despite presenting more isolates, state C had the lowest relative frequency (Fisher’s test p <0.05) of strains classified as virulent and multi-virulent compared to states A and B, as well as multi-virulent profiles (Figure 2).

**Figure 2.**
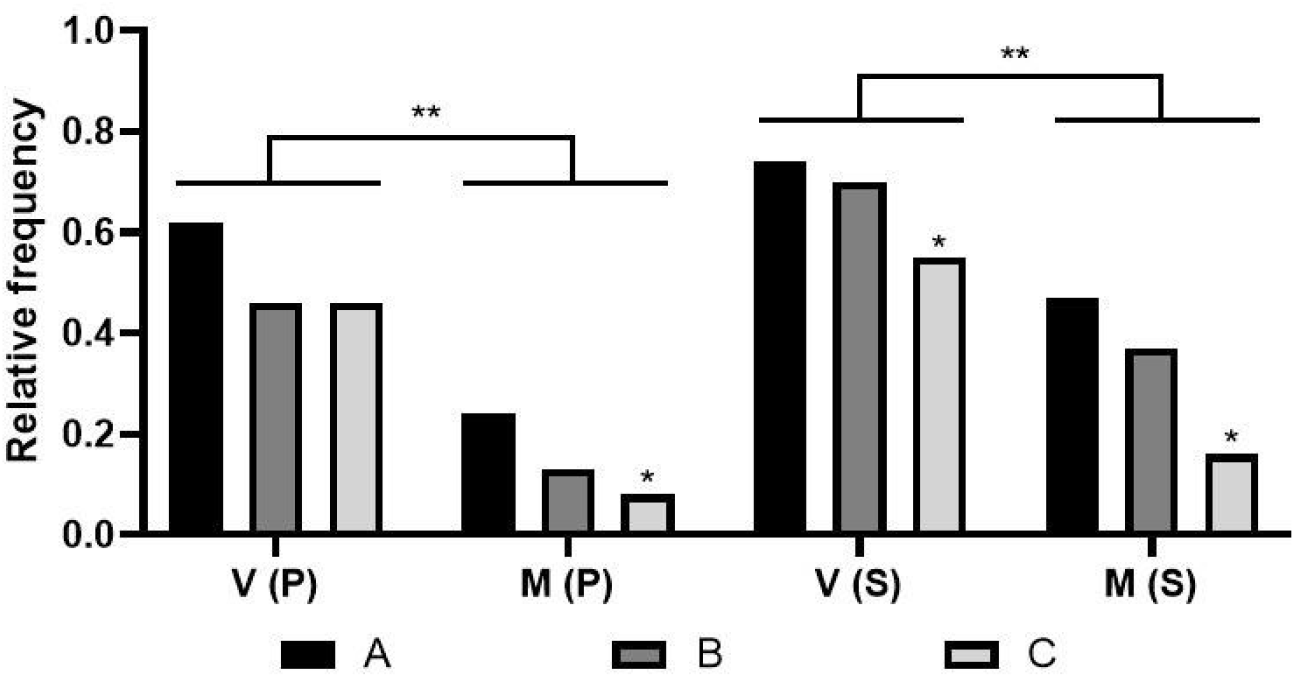
Relative frequency of virulent and multi-virulent strains by state. V: Virulence; M: Multi-virulence; P: profiles; S: strains; States: A, B and C; * p <0.05; ** p <0.005 - One way Anova and Fisher’s test.

The construction of the heat graph led to the identification of more evident hot zones in state A, where we observed maximum expression of relative frequency of genes linked to SGB compared to the other states. Genes linked to biofilm formation were the least frequent, but predominant in all states in the month of December and most common in state A. The highest concentration of virulent and multi-virulent strains was also more expressive in state A, especially in the months of November and December. Thus, state A presents itself as the main hotspot in terms of potential for maintenance of virulent and multi-virulent strains of *C. jejuni* compared to the other states. The other genetic categories (SS, IC and AE) showed a similar frequency in the three states (Figure 3).

**Figure 3.**
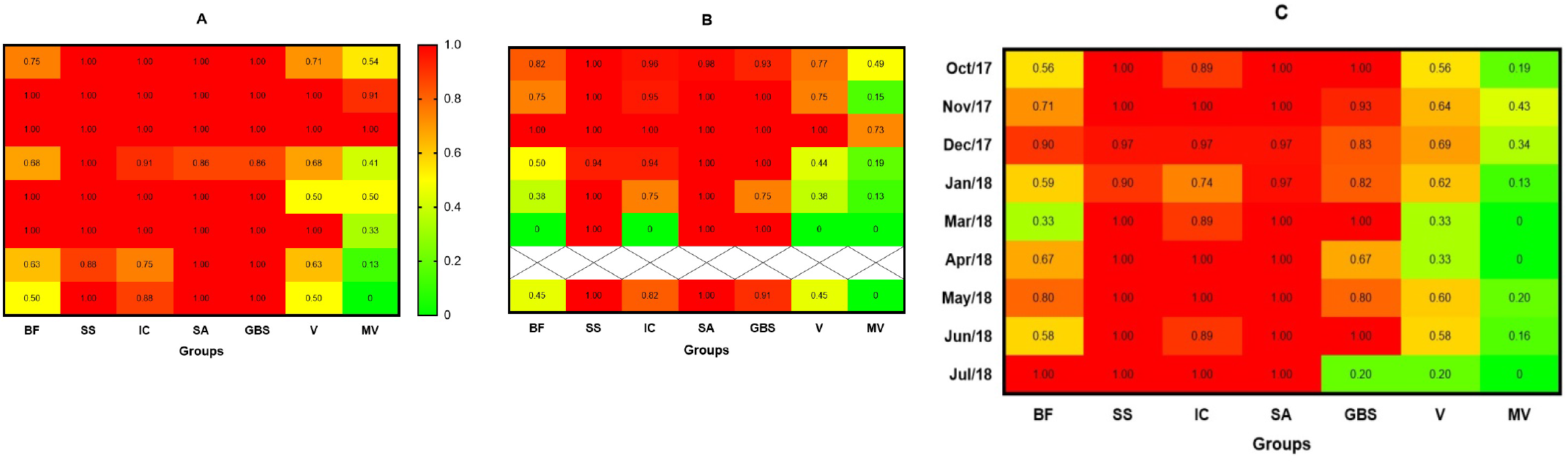
Heat graph defined based on the extreme colors from green to red that indicates the relative frequency of each virulence category studied according to the state and months of isolation of *C. jejuni*. BF: Biofilm formation group (frequency of one or more genes). SS: Secretion system (frequency of one or more genes). IC: Invasion / Colonization (frequency of one or more genes). SA: Stress adaptation (frequency of one or more genes). GBS: Guillain-Barré syndrome (frequency of one or more genes). V: index of virulent strains. MV: index of multi-virulent strains. X: absence of isolates. (GraphPad Prism 8.0.1 software).

Even with a significant number of profiles that contemplate only one strain in the three states, no state had a high I.Var., due to the large number of strains being grouped to a few profiles (Table 6). At the state C the variability below 0.7 is due to the existence of two profiles comprising 16 and 18 strains. In state B, the lower value is linked to a profile that includes 37 multi-virulent strains and in state A to a multi-virulent profile with 25 strains.

**Table 6.**
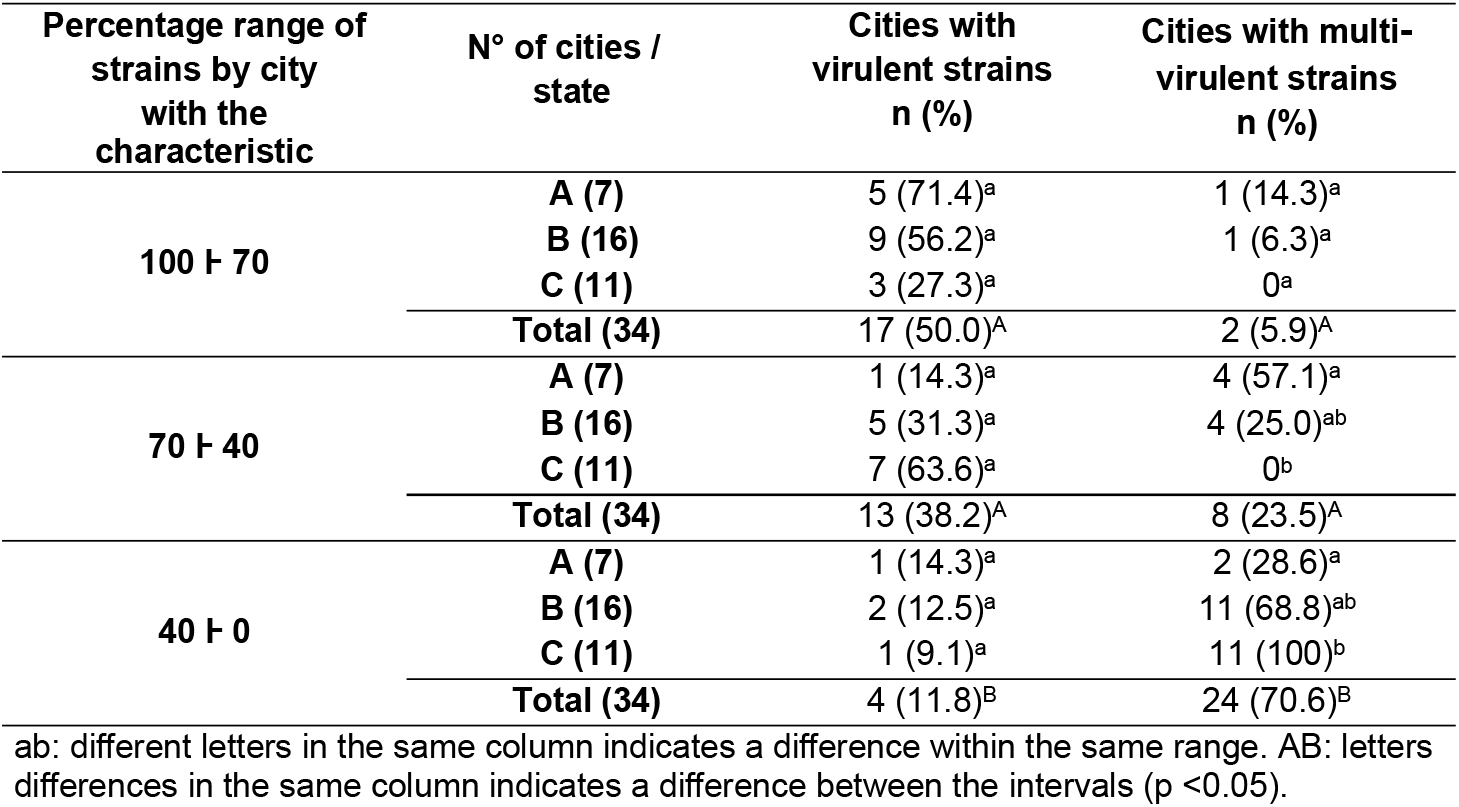
Distribution of cities according to the percentage value of virulent and multi-virulent strains of *C. jejuni* isolated in municipalities located in three Brazilian states (A, B and C).

There was a variation of one to 23 isolates per city, with a median of eight isolates. To determine the I.Var. and the distribution of virulence profiles according to the classification of V and MV, the selected municipalities were the ones that presented three or more isolates, which corresponded to 79.1% (34/43) of the total, being seven in state A, 16 in state B and 11 in state C. The average I. Var. identified in the municipalities was 0.78 and considered high. The significant number of cities with I.Var. greater than 0.7 (23/34 - 70.6%) determined the high mean value identified (Fischer’s test - p <0.05). The number of cities/states with virulent strains did not fluctuate significantly, regardless of the percentage range of virulent strains. However, a significant minority of cities (4/34 - 11.8%) had 40% or less of their virulent strains. Controversially, cities with 0-40% of their multi-virulent strains were predominant (24/34 - 70.6%) in our study. This high number is due to the fact that 100% of the municipalities in state C fall within this range (Table 6).

The I.Var. and the virulence and multi-virulence characteristics were evaluated for the 359 strains, considering the months of October to December of 2017 and January and March to July of 2018.

High I.Var. are directly related to low rates of multi-virulence due to the odds ratio (Table 7). Our study showed that in the months from January to July 2018 we had the lowest rates of multi-virulence, with a decreasing characteristic over time. This determined the high genetic variability between the strains, due to the diversity of virulence profiles. In the months from October to December of 2017, there was greater balance and equivalence in these indexes (Figure 5).

**Table 7.** Frequency of virulence and multi-virulence of strains and profiles by seasonality.

| Month/year | Virulence – n(%) |  | Multi-virulence – n(%) |  | Total |  | OR<br>I.Var. - I.M.<sup>-1</sup> |
| --- | --- | --- | --- | --- | --- | --- | --- |
|  | Strains | Profiles | Strains | Profiles | Strains | Profiles |  |
| **October/2017** | 73 (67.6)<sup>ac,A</sup> | 26 (54.2)<sup>ab,C</sup> | 44 (40.7)<sup>a,B</sup> | 11<br>(22.9)<sup>a,D</sup> | 108 | 48 | 1.16 |
| **November/2017** | 37 (82.2)<sup>ad,A</sup> | 16 (66.7)<sup>a,C</sup> | 20 (44.4)<sup>a,B</sup> | 6 (25.0)<sup>a,D</sup> | 45 | 24 | 1.43 |
| **December/2017** | 37 (78.7)<sup>ad,A</sup> | 13 (61.9)<sup>ab,C</sup> | 21 (44.7)<sup>a,B</sup> | 2 (9.5)<sup>a,D</sup> | 47 | 21 | 1.0 |
| **January/2018** | 41 (53.2)<sup>bc,A</sup> | 15 (37.5)<sup>b,C</sup> | 16 (20.8)<sup>bc,B</sup> | 3 (7.5)<sup>a,D</sup> | 77 | 40 | 4.12 |
| **March/2018** | 8 (42.1)<sup>b,A</sup> | 6 (35.3)<sup>ab,C</sup> | 3 (15.8)<sup>bd,A</sup> | 3 (17.6)<sup>a,C</sup> | 19 | 17 | 45.33 |
| **April/2018** | 4 (57.1)<sup>bcd,A</sup> | 4 (57.1)<sup>ab,C</sup> | 1 (14.3)<sup>ad,A</sup> | 1 (14.3)<sup>a,C</sup> | 7 | 7 | ∞ |
| **May/2018** | 8 (61.5)<sup>bcd,A</sup> | 7 (58.3)<sup>ab,C</sup> | 2 (15.4)<sup>ad,B</sup> | 2 (16.7)<sup>a,C</sup> | 13 | 12 | 66.0 |
| **June/2018** | 20 (52.6)<sup>bc,A</sup> | 15 (62.5)<sup>ab,C</sup> | 2 (5.3)<sup>d,B</sup> | 2 (8.3)<sup>a,D</sup> | 38 | 24 | 30.86 |
| **July/2018** | 1 (20.0)<sup>b,A</sup> | 1 (20.0)<sup>ab,C</sup> | 0<sup>abd,A</sup> | 0<sup>a,C</sup> | 5 | 5 | ∞ |

Different lowercase letters in the same column indicates a significant difference. A B: difference in virulence and multi-virulence of strains (line). C D: difference in virulence and multi-virulence of the profiles (line) (p < 0.05). OR: Odds Ratio, I.Var .: variability index, I.M .: multi-virulence index.

**Table 8:**
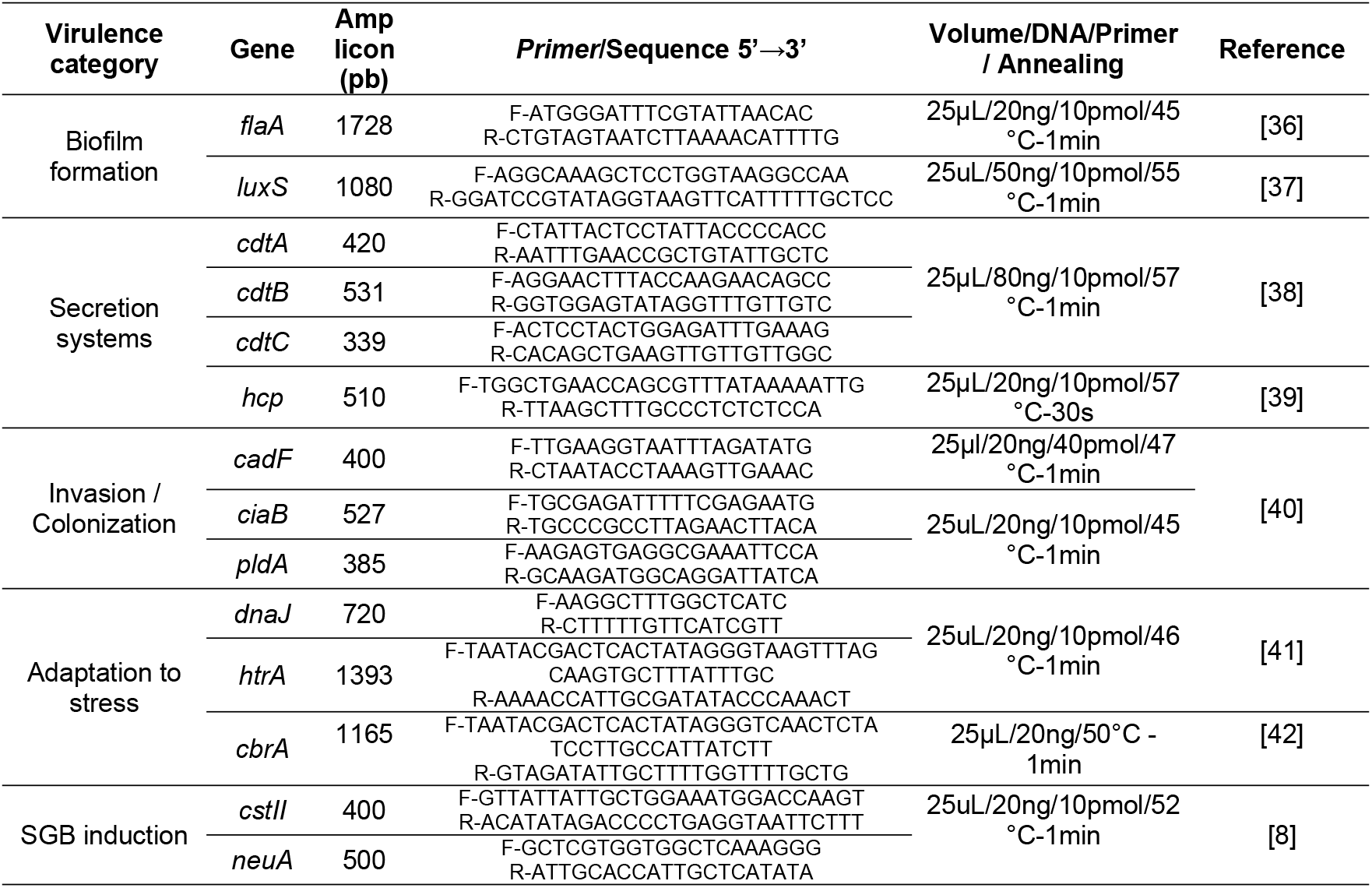
PCR conditions, nucleotide sequences and sizes of amplicons to identify virulence genes in *C. jejuni*.

In general, the months considered hot and humid (Oct-Dec) in the southern hemisphere determine the highest number of isolates associated with high rates of virulence and multi-virulence, indicating the seasonal characteristic of *C. jejuni* (Table 7).

## DISCUSSION

The importance of *C. jejuni* as the main food pathogen, the involvement of chicken meat as a source of infection for humans, and the role of Brazil as an exporter of this food instigate a better knowledge of the molecular and phenotypic epidemiological aspects of these bacteria in the country.

In general, the evolution of bacterial populations is determined by genetic events that permeate the diversification of molecular characteristics. In particular, for *Campylobacter* there is a probable selection pressure due to adaptive characteristics to stress conditions that directly or indirectly influence their suitability, specifically, in species adapted to the host, but widely exposed to hostile environmental conditions, as is the case of *C. jejuni* [11]. The significant prevalence of strains that concomitantly present genes associated with adaptation to stress (309/359 - 86.07%) (Table 2) suggests the existence of strains capable of survival in several niches. In parallel, this adaptive potential can facilitate its dissemination, since there is an intensification of the survival rate under thermal, osmotic, and oxidative stress. Especially for *dnaJ*, which encodes heat shock proteins, particularly important for the survival of *Campylobacter* in foods that are kept during marketing under refrigeration or freezing, as well as during their preparation, if they are undercooked [12].

The high prevalence of the presence of the *hcp* gene (97.77%) linked to the category of secretion systems, indicates the potential to cause more serious infections associated with bloody diarrhea [13], since it is involved in the type VI secretion system (T6SS), responsible for carrying toxins into the environment, prokaryotic [14] or eukaryotic [15] cells. The other genes related to the secretion system (*cdtABC*) had lower and similar frequencies (42 to 48%) among them in our study.

In general, the complete presence of the CDT genetic complex indicates the potential for release of a functional cytotoxin that contributes to virulence, invasion, and cell adhesion and is essential for the release of interleukin-8 (IL-8) that contributes to the inflammatory response of the host mucosa, besides favoring the loss of the integrity of the intestinal epithelium, of the cellular junctions and inducing apoptosis.

The significant difference in the prevalence of genes linked to the CDT complex and the *hcp* gene may be related to the fact that, although they are important secretion systems in the pathogenesis of *C. jejuni*, both are encoded in independent regions of the genome, and therefore, are closely related [16,17]. Similar to the CDT complex, genes linked to the invasion/colonization category had a concomitant prevalence lower than expected (63.79%) when compared to other studies conducted in Brazil [18, 19]. Some studies point out that the limitation in the pathogenic potential, in particular to the initial infection processes (invasion/colonization), observed in some strains of *Campylobacter*, presents benefits to the pathogen, due to the easier escape of the immune system combined with maintenance in the host in the long term through the establishment of chronic infection [20,21].

It is estimated that 25 to 50% of cases of GBS can be associated with previous campylobacteriosis [22]. The presence of the *neuA* and *cstII* genes demonstrated a high relationship with the production of lipopolysaccharides (LOS) from the *Campylobacter* cell wall involved in the development of the syndrome [7, 23] and identified in 88.86% and 77.44% of the strains, respectively in our study. These values are alarming due to their superiority compared to the findings in the literature [7, 8]. However, despite a large number of isolates with a potential risk of developing post-infection neuropathy, the host’s immune system (humoral and cellular immunity) is primarily responsible for the development of GBS.

The joint prevalence of genes related to the biofilm formation/motility virulence category was 182/359 (50.7%). The association of both characteristics in our study was due to the physiological involvement identified for the *luxS* and *flaA* genes in *C. jejuni* associated with the complexity of the biofilm formation process that can be reduced by up to 57% in strains that do not share both genetic factors. [24]. The high capacity to form biofilms in production environments is quite evident for *C. jejuni* and involves several intrinsic and extrinsic mechanisms [7]. But, especially for our strains that do not share the biofilm/motility category (177/359 - 49.3%), the production of biomass can be significantly compromised [24] which can facilitate the control of these pathogens by the hygiene processes used by industry.

Considering the clusters identified in the 124 profiles, the significance of cluster A16, which includes virulent (57) and multi-virulent (14/57) profiles, is evident (Table 2). This fact sets up the discussion aimed at the virulence and multi-virulence indices identified from different perspectives.

We observed a similarity in the number of establishments and municipalities that presented more than 70% of their strains classified as virulent (21/46 and 17/43, respectively), of which in two cities/establishments these strains were multi-virulent, significant values compared to the lowest interval (Tables 4 and 6). In a broader analysis, it became clear that these locations do not include state C in a significant way (Figure 2), but mainly the state of A (Figures 3), whose strains had a more dispersed origin in the territory, but with a convergence of the indices of virulence and multi-virulence to the central and metropolitan regions of the state. It is likely that this state concentrates the largest arsenal of *C. jejuni* with a higher evolutionary level in comparison to the other states involved in the study and, therefore, it is the main hotspot for the maintenance of virulent strains since over time this species tends to acquire a greater number of virulence genes [19]. There is also a mention that the inclusion of control measures and legislation in establishments strengthens a reduction in the percentage of isolation while promoting a pressure to select more virulent strains since the obstacles in environmental selection favors the preservation of more competitive strains [25]. This fact was also identified in state A, considering the restricted and significant number of strains isolated compared to the other states.

At the temporal level, it is evident that the quantitative distribution (Table 7), the virulent and, mainly, the multi-virulent character of the strains (Figure 4) present a seasonal pattern so that in the hottest and humid months (October to December) of the year we identify the higher rates. This behavior was also recorded in a study carried out on *C. jejuni* isolated from chicken carcasses that showed a higher prevalence and virulence in the summer compared to the winter [26]. The analysis of the variability indices showed discrepancies in the values found in the general analysis of the data and at the state level (less than 0.7) with the values obtained by sample, by the establishment, and by the municipality (greater than 0.7). The high variability between strains isolated from the same sample (I.Var. = 0.82) indicates the existence of different genotypes of *C. jejuni* coexisting within the same host organism. In fact, this type of adaptation to sub-niches or niches common in the same host is proposed in *C. jejuni*, where up to 10 different genotypes have already been reported in a single chicken carcass [27]. This shows that different strains, especially emerging ones [11], with different degrees of complexity and virulence, can cohabit the same host in a commensal manner. In parallel, the similarity in I.Var. found in the strains considering establishments and municipalities (average greater than 0.7) was expected. This is due to the fact that in only three of the 43 investigated cities, we had two different establishments present, for the remainder the number of strains per municipality and per establishment was equivalent. Because it is the same establishment/municipality in most cases, the high variability may be related to the environmental pressure that tends to select different profiles, due to the different managements and conducts adopted in each establishment, and in general, considering the intrinsic differences of the productive process itself, which involve stages in different degrees of stress that are tolerable or not to the different strains [28].

**Figure 4.**
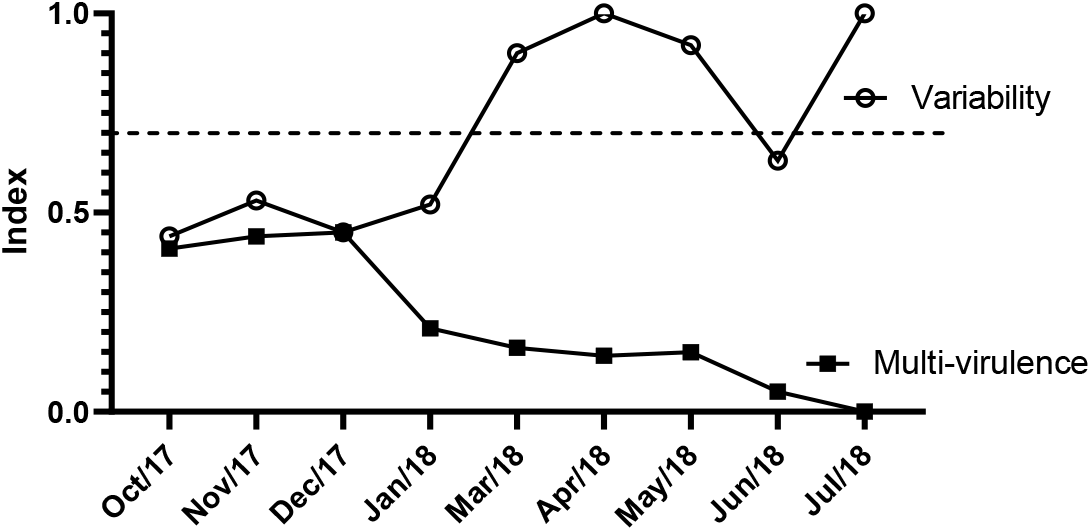
Indexes of variability and multi-virulence of *C. jejuni* over time of isolation. Dashed line (---) separates low and medium indexes from values of high genetic variability.

At the same time, *C. jejuni* has high genetic plasticity due to its high potential to carry out horizontal gene transfer and recombination, since the bacterium is naturally competent for uptake and transformation of DNA, which favors the diversity of virulence profiles [29]. Paradoxically, the analysis of I.Var. in the total sampling and its distribution at the state level it shows lower values, due to the concentration of strains in the same group (total or by state). These lower values occurred due to the existence of a large number of strains grouped in a few profiles classified as virulent and multi-virulent, which represents an evolutionary trend in *C. jejuni* [19]. The distribution of I.Var. over time showed an inverse association in relation to the multi-virulence index (odds ratio> 1.0) (Table 7). This fact is configured in elevated I.Var. in months with low rates of MV strains. As well as, the occurrence of low variability in the months that includes the summer season, which had the highest rates of MV strains. The steep drop in I.Var. associated with multi-virulence of strains in hot and humid periods of the year is consistent with the significant increases recorded worldwide in enteritis caused by *Campylobacter* associated with a surprising increase in the severity of cases [30]. Considering the complexity of the epidemiology of *C. jejuni*, our results may contribute with strategies for its control. The gene profile demonstrates that the studied strains have the potential for high adaptation to hostile environmental situations, but on the other hand, they have a lower frequency of genes linked to the biofilm formation, indicating that adequate hygiene processes can be strategic for their control. Finally, the high levels of virulence, especially in summer and in state A, suggest the need to adopt control measures converging to these findings.

## METHODOLOGY

### Identification, MALDI-TOF MS

Overall, 359 strains of *C. jejuni* isolated from 114 refrigerated chicken carcasses, slaughtered in establishments registered with the Federal Inspection Service by the Ministry of Agriculture, Livestock, and Supply (MAPA), were analyzed. The carcasses were analyzed from October 2017 to July 2018 to meet the Exploratory Program for the Research and Estimation of the Prevalence of Campylobacter spp., the same ones used for the official verification of the *Salmonella* spp. Control and Monitoring Program, determined by IN nº 20, of October 21, 2016 [34]. The carcasses collections were carried out in establishments located in states A, B, and C, distributed in 43 municipalities.

The strains were isolated and identified as *C. jejuni* in official MAPA laboratories using the methodology proposed by ISO 10272-1: 2017 [35] and the mass spectrometer (Maldi TOF®) for the identification of the genus and species, respectively. Additional information such as establishment, place, and date of isolation were received together with the cryopreserved strains and deposited in the culture bank of the Molecular Epidemiology Laboratory of the Faculty of Veterinary Medicine of the Federal University of Uberlândia (LEPIMOL-FAMEV-UFU).

The samples were prepared according to the direct transfer protocol (Direct Transfer Method) for the plate from the isolated colony in three spots. After application to the plate, the sample was covered with 1 μL of alpha-cyano-4-hydroxycinnamic acid (HCCA) solution. Reference strains belonging to LANAGRO-RS were used as controls. The Brucker platform, model Autoflex Speed, was used to identify the isolates. The spectrum bank was the MALDI Biotyper RTC / OC 3.1. The species decision criteria followed: equal to or greater than 1.7 for genus and equal to or greater than 2.0 for genus and species. The data acquisition control program was Brucker, flexControl 3.4. Calibration was performed using the Protein Standard I calibrator, with a calibration error tolerance of less than 200ppm.

#### Virulence panel

After reactivation and confirmation of the strains by the oxidase, catalase, motility, and typical morphology tests [35], the genomic DNA was extracted using the Wizard Genomic DNA Purification Kit (Promega), following the protocol provided by the manufacturer. The purified DNA (10 ng) was used as a template for all PCR reactions. The PCR conditions and primers used in this study are described in Table 1.

Overall, 14 genes were studied, divided into five virulence categories: i) biofilm formation - *flaA* (motility) and *luxS* (quorum-sensing mechanism); ii) secretion systems - *cdtABC* (distensive cytotoxin secretion) and *hcp* (type VI secretion system; iii) invasion and colonization - *cadF* (intracellular colonization), *ciaB* (cell invasion) and *pldA* (invasion / colonization); iv) adaptation to stress - *dnaJ* (thermotolerance), *htrA* (aid in growth under stress) and *cbrA* (resistance to osmotic shock); and v) induction of Guillain-Barré Syndrome - *cstII* (SGB) and *neuA* (SGB).

PCR reactions were performed using the GoTaq® Green Master Mix kit (Promega), according to the manufacturer’s instructions. As positive controls, strains of *C. jejuni* ATCC 33291, IAL 2383, and NCTC 11351. The amplified products were subjected to electrophoresis in 1.5% agarose gel, using the TBE 0.5x running buffer (Invitrogen) and as a molecular weight standard, the 100bp marker (Invitrogen).

For the classification of the virulence level of the strains, a criterion adapted from the analysis of the antimicrobial multidrug resistance index was used [43] and the five categories described in Table 1. Thus, the strains were classified as virulent (V) when they had at least one gene linked to each category and multi-virulent (MV) when they had two or more genes from each of the five categories.

The variability of the strains within a given group (sample, establishment, municipality, state) was verified using the analysis of the number of distinct virulence profiles identified. The variability, virulence, and multi-virulence indices were determined by calculating the relative frequency for each group of strains evaluated. The classification for the variability index was adapted [43], being low for values less than 0.5, medium for values between 0.5 and 0.7, and high for values greater than 0.7.

### Statistical analysis

The analysis of the results of the virulence panel was performed based on the number of strains isolated in each analyzed sample, in the establishment of origin, in the place (city and state), and in the seasonal aspects for the percentage description. For comparative analyzes, the normality of the data was verified, followed by the application of Fischer’s exact test or Student’s t-test to compare two variables and ANOVA or Kruskal-Wallis in the comparison of three or more variables. The program used was Graph Pad Prism 8.0.1 with a 95% of a confidence interval.

## ACKNOWLEDGMENTS

This study was financed in part by the Coordination for the Improvement of Higher Education Personnel – Brazil (CAPES) – Finance Code 001

## AUTHOR CONTRIBUTIONS

Conceived and designed the experiments: PABMP, DAP. Performed the experiments: PABMP, ECAL, ASMB, MdVF, ABGB, EPM. Analyzed the data: PABMP, RTdM, ALG, GPM. Contributed reagents/materials/analysis tools: DAR, PMA, FB, TFP. Wrote the paper: PABMP.

